# Study on Early pregnancy diagnosis in sows based on joint analysis of vaginal secretion metabolomics and machine learning

**DOI:** 10.1101/2025.09.25.678563

**Authors:** Yun Feng, Ruonan Gao, Yun Zhang, Wengang Yang, Mengxun Li, Qingchun Li, Guang Pu, Yongsheng Zhang, Yujun Ren, Zikai Ai, Kun Yan, Huiwen Lu, Tao Huang

## Abstract

**Objective:** Metabolic transformations in female mammals throughout gestation manifest in the metabolite profiles of bodily fluids, potentially serving as biomarkers for the diagnosis of pregnancy. This investigation concentrated on sows 18 days post-mating, with the objective of identifying biomarkers pertinent to early pregnancy diagnosis through the analysis of variations in metabolic molecules within vaginal secretions.

**Methods:** Vaginal secretion samples were procured from both pregnant and non-pregnant sows 18 days post-mating, subsequently subjected to untargeted metabolomic analysis utilizing liquid chromatography-mass spectrometry (LC-MS). The samples were partitioned into training and testing datasets, and machine learning algorithms—specifically, Random Forest (RF) and support vector machine (SVM)—were integrated with metabolite weight ranking to identify key biological markers.

**Results:** A total of 3,249 metabolic molecules were identified, among which 534 exhibited significant differential expression in the vaginal secretions of pregnant versus non-pregnant sows. Notably, three hormones that correlate with the pregnant status of the sow were discerned: Progesterone, 3-Deoxyestradiol, and Prostaglandin E1. KEGG pathway analysis revealed that the differentially expressed metabolites were predominantly enriched in nucleotide metabolic pathways. The RF model demonstrated an impressive accuracy of 1.00 in the training dataset and 0.88 in the testing dataset, while the SVM model maintained an accuracy of 1.00 across both datasets. Utilizing model weights and differential expression data, three pivotal metabolites were identified: Indolepropionic acid, cis-Aconitate, and 4-p-Coumaroylquinic acid, which exhibited ROC curve AUC values of 0.90, 0.89, and 0.79, respectively.

**Conclusion:** This study validates the viability of utilizing vaginal secretions for the early diagnosis of pregnancy in sows, thereby establishing a foundation for the advancement of non-invasive diagnostic methodologies.

## INTRODUCTION

In the pig industry, the augmentation of the number of piglets weaned per sow per annum (PSY) through the optimization of sow breeding practices remains a paramount objective. However, in practical production scenarios, the failure of sow pregnancies attributable to a myriad of factors extends non-productive periods, thereby leading to considerable economic detriment for pig farms [1]. Consequently, the timely assessment of sow gestational status, coupled with the swift remating of non-pregnant sows, constitutes essential strategies to mitigate non-productive days and enhance sow PSY.

At present, the predominant method for diagnosing pregnancy on pig farms is ultrasonic detection, which utilizes ultrasound technology to discern physiological alterations in the amniotic fluid or fetal development within the uterus of the sow [2,3]. However, this technique produces reliable results only after a period of 22 to 25 days post-mating, while the average estrous cycle of the sow encompasses merely 21 days. This temporal discrepancy inherently prolongs the duration of non-productive days associated with unsuccessful pregnancies, thereby exacerbating economic losses. Alternative diagnostic approaches—such as boar estrus testing, platelet counting, radioimmunoassay, rectal palpation, and biohormonal analysis—are hampered by significant constraints, including heightened operational complexity, increased costs, or considerable stress imposed on the sows [4,5].

In recent years, investigations into vaginal secretions have garnered increasing attention, particularly concerning their application in the diagnosis of reproductive tract disorders in female mammals. Nonetheless, there exists a dearth of studies examining the relationship between the physicochemical characteristics of vaginal secretions and the status of pregnancy. Metabolomics is a specialized field devoted to the quantitative examination of animal metabolites and the exploration of metabolic transformations induced by various physiological conditions. The foundational premise of this discipline posits that physiological alterations within organisms precipitate concomitant changes in the chemical composition of metabolites. Consequently, metabolomics has been widely employed to investigate transitions between different physiological states in living organisms [6–8].

Machine learning, a crucial domain within the realm of artificial intelligence, constitutes the fundamental basis for the advancement and actualization of AI technologies. It exhibits remarkable proficiency in comprehending and synthesizing multifaceted data, and when synergistically integrated with histological studies, it facilitates the effective extraction and assimilation of extensive, intricate datasets across a plethora of histological contexts. As a result, machine learning has ascended to prominence as a formidable research instrument, producing promising advancements across numerous scientific disciplines [9–11].

In this research, we utilized liquid chromatography-mass spectrometry (LC-MS) in conjunction with advanced machine learning algorithms, specifically Random Forest (RF) and Support Vector Machine (SVM), to elucidate the metabolomic profiles of vaginal secretions from non-pregnant and pregnant sows. Our primary objectives were to discern differentially expressed metabolic constituents in vaginal secretions across the two cohorts and to investigate the potential of employing these vaginal secretions, augmented by machine learning models, as a robust means for early pregnancy diagnosis in sows.

## MATERIALS AND METHODS

### Experimental material Test animal handling

All experimental samples utilized in this study were generously supplied by Xinjiang Tiankang Animal Husbandry Science and Technology Co., Ltd. The subjects comprised adult binary replacement sows, ranging in age from 225 to 250 days and weighing between 135 and 150 kg, all exhibiting excellent physical health. These animals were kept under uniform feeding and housing conditions: the sows presented no evidence of genetic disorders or physical deformities and were maintained in an environment with temperatures between 20 and 25°C and humidity levels of 50 to 55%. The production system adhered to an all-in/all-out strategy with the provision of a nutritionally complete feed. Following synchronized estrus, the sows underwent batch mating, and successful pregnancies were confirmed through ultrasonography performed 23 to 25 days post-mating.

### Sample collection

Eighteen days following batch mating, a selection of fifteen sows was made at random. A sterile, long-handled medical swab was carefully inserted 10–20 cm into the vaginal canal of each sow, retained for a duration of 30 minutes, and subsequently withdrawn. Simultaneously, vaginal secretion samples were procured from ten additional sows, also randomly selected from the same batch, which had either failed to conceive or were not mated. All samples were promptly frozen in liquid nitrogen and thereafter stored at a temperature of −80°C.

### Non-targeted metabolomics analysis

#### Sample Preparation

The samples were initially thawed at a temperature of 4°C and subsequently subjected to sonication in an ice bath for a duration of 30 minutes. All samples were then transferred to new Eppendorf tubes and subjected to lyophilization. Following this, 200 μL of pre-cooled water and 800 μL of pre-cooled methanol were introduced, followed by vigorous vortexing to ensure thorough mixing. The samples underwent another round of sonication in an ice bath for 20 minutes, after which protein precipitation was achieved through incubation at −20°C for 1 hour.

Centrifugation was performed at 16,000g for 20 minutes at 4°C, and the resulting supernatant was carefully collected. Upon air-drying the precipitated proteins, 200 μL of SSDT (Sodium Dodecyl Sulfate sample buffer) was added to enable BCA (Bicinchoninic Acid) quantification of the proteins, with the resulting quantitative protein concentrations depicted in Fig. 1.

**Fig. 1.**
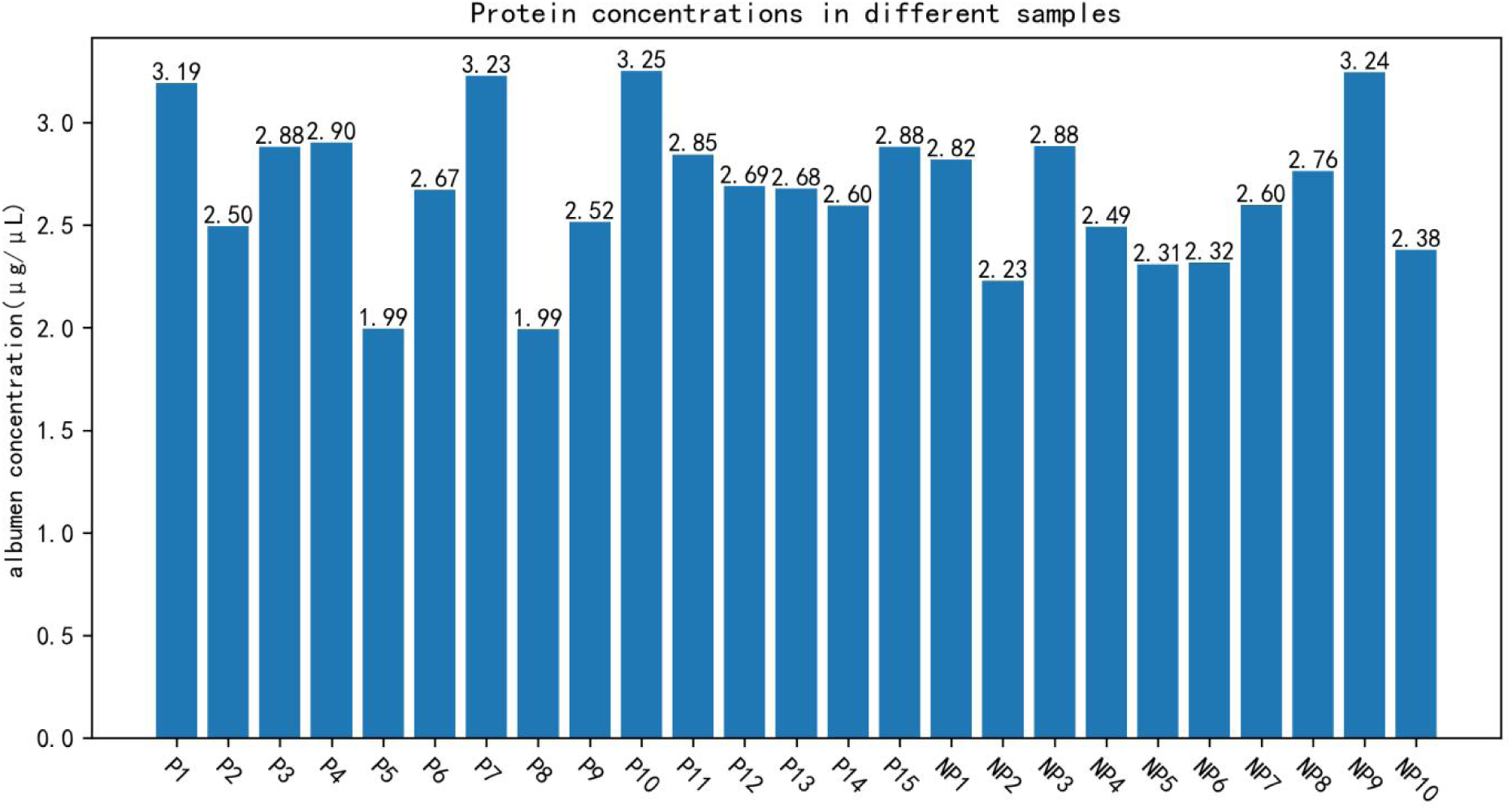
Protein concentration in different samples.

Based on the protein content quantified in the samples, the corresponding cellular concentrations across the different samples were deduced. Subsequently, supernatants containing equal cell concentrations from each sample were collected and subjected to drying using a high-speed vacuum concentration centrifuge. For mass spectrometry analysis, 35 μL of a pre-cooled methanol-water solution (1:1, v/v) was prepared.

#### Liquid Chromatography Mass Spectrometry

The samples were maintained in an autosampler at a temperature of 4°C throughout the duration of the analysis. The analytical procedures were conducted utilizing a Shimadzu Nexera X2 ultra-high-performance liquid chromatography (UHPLC) system, equipped with an ACQUITY UPLC® HSS T3 column (2.1 × 100 mm, 1.8 µm) (Waters, Milford, MA, USA). An injection volume of 4 μL was employed, with the column temperature set to 40 °C and a flow rate of 300 μL/min.

The mobile phases for the chromatographic separation comprised Phase A: an aqueous solution of 0.1% formic acid, and Phase B: a 0.1% formic acid solution in acetonitrile. The chromatographic gradient elution program was meticulously delineated as follows: from 0 to 1 minute, Phase B was held constant at 2%; from 1 to 5 minutes, Phase B underwent a linear transition from 2% to 48%; from 5 to 7 minutes, Phase B progressed linearly from 48% to 80%; from 7 to 11 minutes, Phase B sustained at 80%; and from 11 to 13 minutes, Phase B increased linearly from 80% to 100%. Subsequently, from 13 minutes to 13.01 minutes, a linear variation occurred.

Subsequently, each sample was analyzed using electrospray ionization (ESI) in both positive (+) and negative (-) ion modes. The mass spectrometry analysis was conducted with a TripleTOF® 6600 mass spectrometer (ABSCIEX) operating in information-dependent acquisition (IDA) mode for both ionization polarities.

### Statistical analysis

In this study, following the normalization of mass spectrometry ion peak intensities (z-score), a comprehensive analysis was conducted utilizing various models. Initially, a dimensionality reduction technique was employed to effectively characterize the data derived from the two distinct sample types. Subsequently, the data underwent fitting and screening processes to identify significant characteristic metabolic molecules through the application of machine learning models.

### Lower dimensionality

The principal methodologies for dimensionality reduction utilized in this investigation were principal component analysis (PCA) and partial least squares discriminant analysis (PLS-DA). PCA is an unsupervised dimensionality reduction technique that transforms high-dimensional data into a more manageable low-dimensional space by discerning principal components—orthogonal directions that encapsulate the maximum variance in the data. This process ensures the utmost retention of the original informational content. Conversely, PLS-DA, a supervised learning framework, elucidates predictive relationships between two datasets by modeling their covariance structures. This approach proves particularly efficacious when addressing high-dimensional variables within the constraints of small sample sizes. Furthermore, volcano plots were employed to illustrate P-values and fold changes in the context of differential expression analysis.

### Machine Learning Classifiers

Subsequent to the acquisition of mass spectrometry data, the dataset was judiciously partitioned into training and test subsets in a ratio of 7:3. Subsequently, the RF and SVM models were trained on the training dataset, and the efficacy of each model was assessed utilizing the test dataset following the completion of the training process.

The principal mechanism underlying the application of Mean Decrease Impurity (MDI) to evaluate the significance of model features within the RF framework is predicated on assessing the mean decrement of Gini Impurity (GI) attributable to a given feature throughout all constituent trees. This process commences with the calculation of the disparity (VIMGini,j,m) between the Gini coefficient of the parent node (GIm) and the weighted mean of the Gini coefficients of its left and right child nodes (GIl, GIr) for every splitting node. Subsequently, the impurity reductions associated with the feature are aggregated across all splitting nodes within an individual tree (VIMGini,i,j). The culmination of these values is then averaged across all trees in the ensemble to yield the overall importance metrics for the feature in question (VIMGini,j). The formula is as follows:

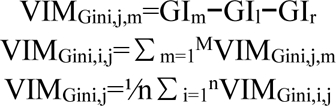

In the linear kernel SVM framework, the decision function is articulated as f(x)=**w***^T^***x**+*b*, wherein **w** signifies the weight vector, *b* represents the bias term, and **x** denotes the feature vector. The coefficients of the weight vector serve as direct indicators of the influence exerted by each feature on the classification decision. Consequently, the significance of the features within the model can be quantified through the absolute values of these weight coefficients.

In this study, the RF model was configured with 100 estimators (n_estimators = 100) and applied default hyperparameters, whereas the SVM model utilized a linear kernel function with a regularization parameter (C) set to 1, maintaining the default settings for all other hyperparameters.

Metabolites exhibiting substantial and pronounced differences were identified through a comprehensive expression significance analysis, alongside RF and SVM methodologies. Subsequently, the disparities in their expression levels were meticulously quantified, following which ROC curves were generated. Corresponding metrics, encompassing AUC values, optimal thresholds, sensitivity, specificity, and Youden’s index, were computed for each ROC curve.

## RESULTS

### Comparison of protein concentration among samples

As illustrated in Figure 1, the concentrations of proteins exhibited significant variation across the vaginal secretion samples, mirroring the corresponding fluctuations in cellular density. Subsequently, the samples were standardized to ensure uniform cell quantities for subsequent analyses, predicated on the quantification of their protein concentrations.

### QC sample mass spectrometry total ion flow results confirm LC-MS reliability

Figure 2 presents the correlation analysis graphs derived from the QC samples. The trio of scatter plots situated in the lower left quadrant illustrates the expression levels of various metabolic molecules across distinct samples. The histograms along the diagonal depict the distribution of data within each sample, while the correlation coefficients for the corresponding pairs of samples are displayed in the three graphs located in the upper right corner. The observations depicted in the graphs reveal that the expressions of various metabolic molecules across different samples exhibit a near-linear relationship. Furthermore, the expression data for each metabolic molecule approximates a normal distribution. The correlation coefficients among the samples exceed 0.9, indicating a remarkably high degree of concordance. This consistency among the measured data substantiates the reliability of the mass spectrometry results.

**Fig. 2.**
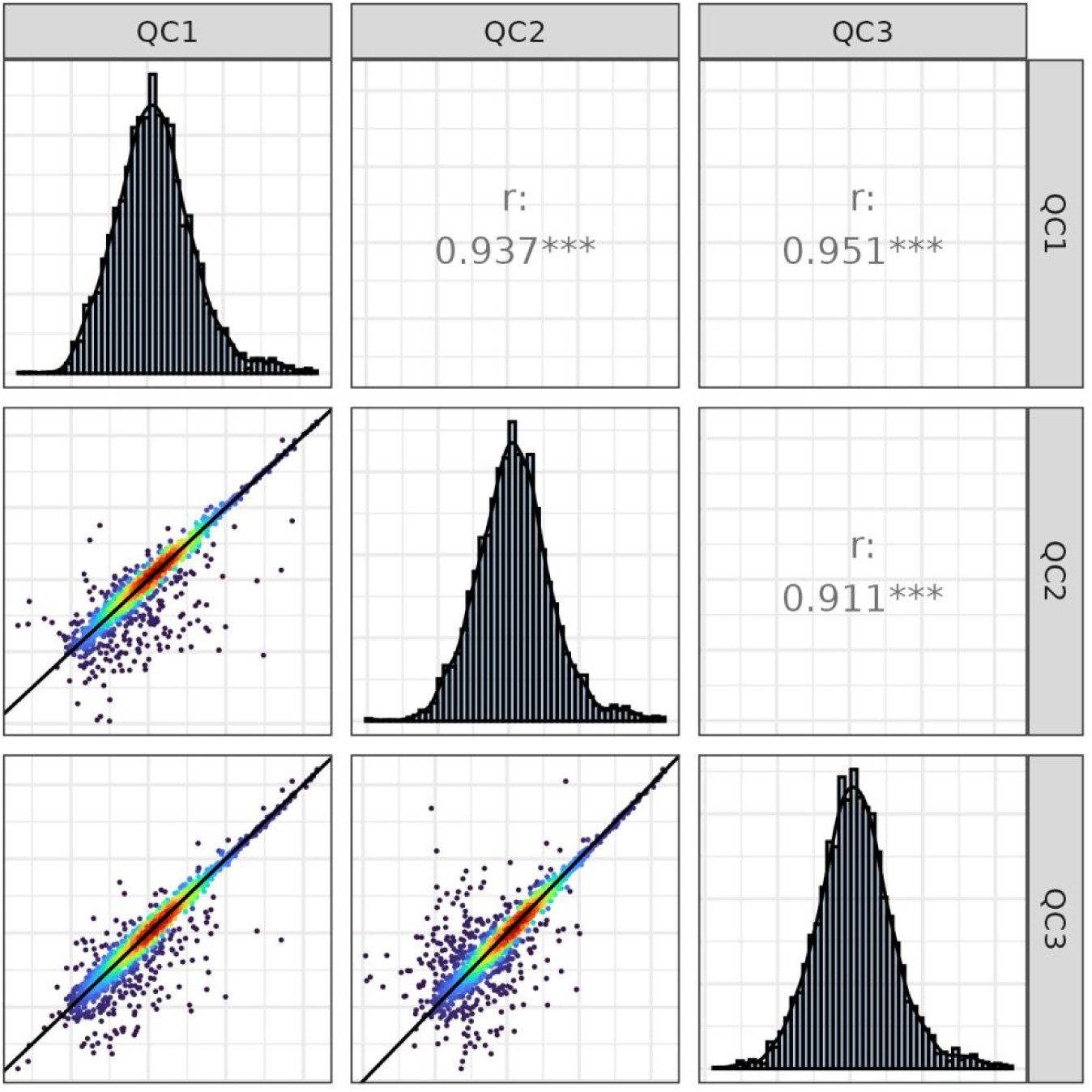
Correlation analysis of quality control samples.

**Fig. 3.**
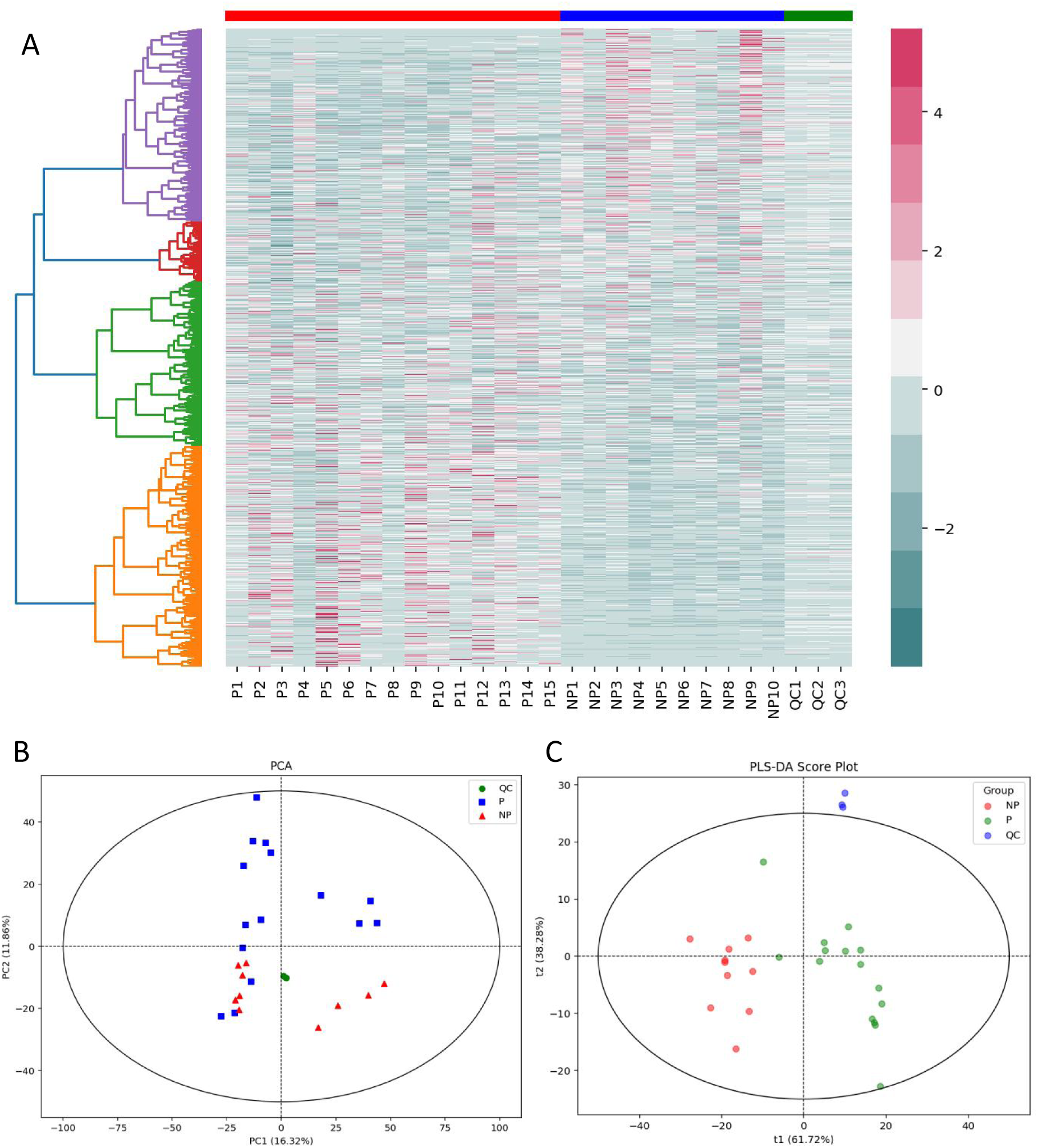
Sample downscaling analysis. (A) Clustering heatmap of pregnant, non-pregnant and QC samples. (B) PCA analysis plot of pregnancy, non-pregnancy and QC samples. (C) PLS-DA analysis plot of pregnancy, non-pregnancy and QC samples.

**Fig. 4.**
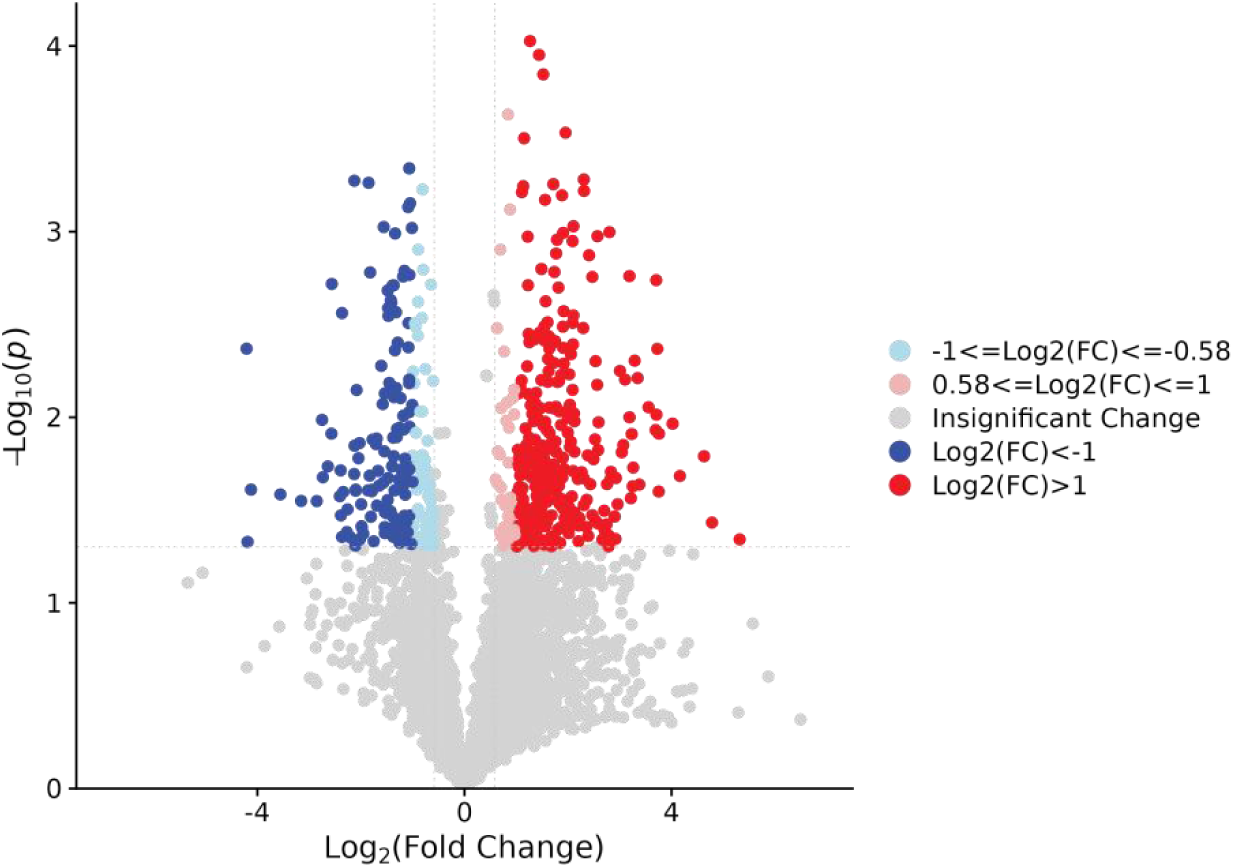
Volcano map of differential metabolites.

### Non-targeted metabolomics assays confirm subgroup sample differences

The hierarchical clustering heat map reveals that the disparities in metabolic molecule expression between pregnant and non-pregnant samples are pronounced. In contrast, the expression levels of each molecule in the QC sample exhibit remarkable consistency, thereby underscoring the stability of the QC sample and affirming the reliability of the experimental findings.

The PCA downscaling analysis of the samples reveals a notable concentration and clustering of the quality control samples within the PCA plot. This observation underscores the stability and reliability of the detection instrument throughout the sequencing process, thereby enhancing the credibility of the obtained results. The graph illustrates a discernible trend of separation between pregnant and non-pregnant samples overall; however, certain pregnant samples are not readily distinguishable from their non-pregnant counterparts, potentially leading to misclassification. Utilizing partial least squares discriminant analysis, a pronounced differentiation trend emerges between samples from pregnant and non-pregnant subjects. This observation underscores the substantial disparities in the metabolic characteristics of vaginal secretions between pregnant and non-pregnant sows. Furthermore, the quality control samples exhibit a centralized distribution within the graphical representation, thereby affirming the robustness and reliability of the test results.

In this investigation, we identified a total of 3,245 distinct metabolic molecules within the vaginal secretions of sows. Of these, 510 metabolic molecules exhibited significant differential expression between pregnant and non-pregnant sows. Notably, 323 of these molecules were markedly up-regulated, while 187 displayed a significant down-regulation in expression.

Through KEGG enrichment analysis, the metabolic pathways of molecules that were significantly enriched in the top 30 of differential significance within vaginal secretions were identified, as illustrated in Fig. 5. The bubble diagram of the significant pathways reveals that nucleotide metabolism emerged as the predominant pathway, while the most significantly enriched molecules fell under the category of metabolic pathways. In the second-tier KEGG pathway category bubble diagram, nucleotide metabolism continued to stand out as the most significant pathway, alongside other notable pathways including amino acid metabolism and transmembrane transport.

**Fig. 5.**
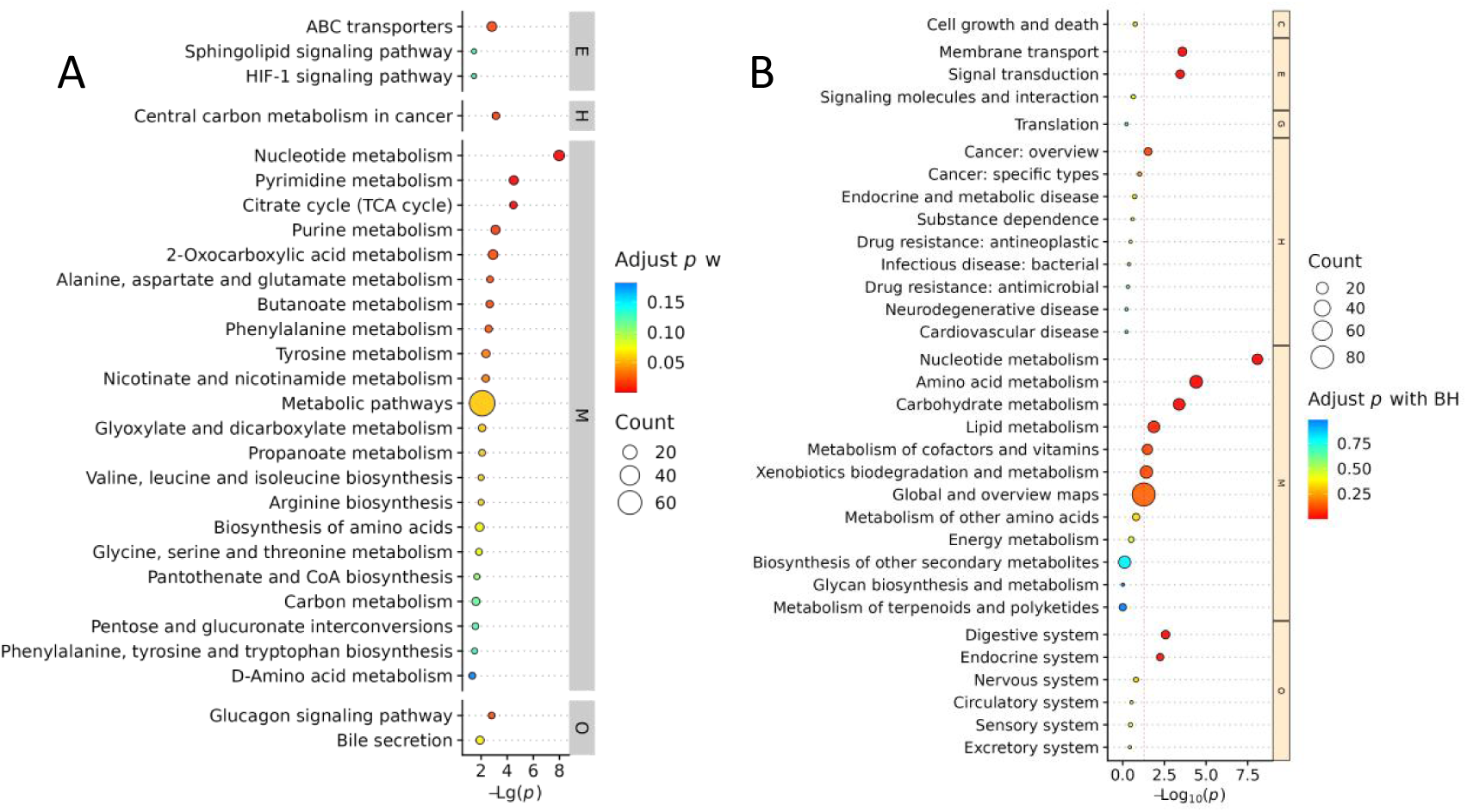
KEGG-enriched pathway analysis. (A) KEGG significant pathway category analysis. (B) Bubble diagram of KEGG pathway categories in the second layer.

**Fig. 6.**
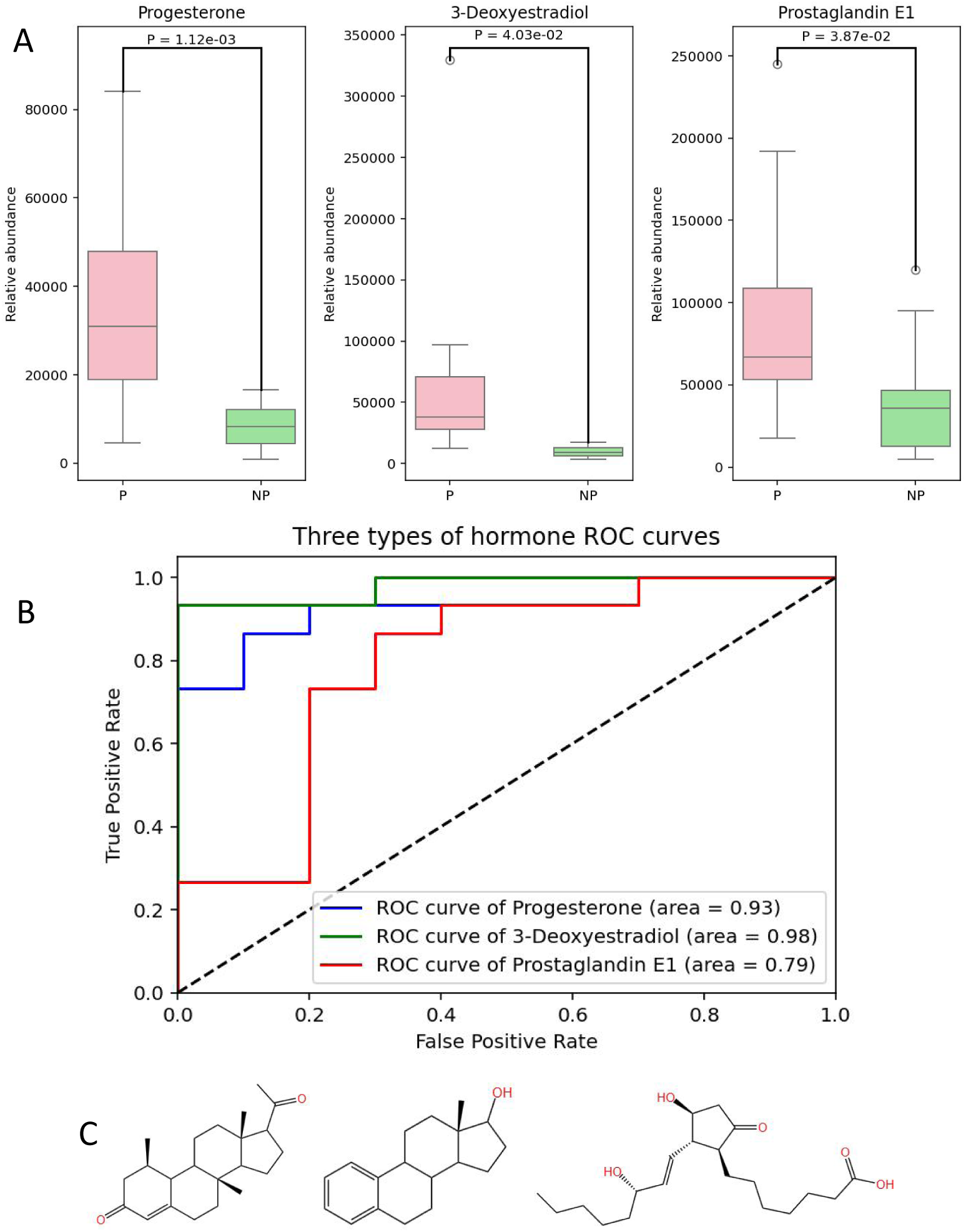
Three pregnancy-related hormones in vaginal secretions. (A) Box line plot of the expression of three pregnancy-related hormone molecules. (B) ROC curves of three pregnancy-related hormone molecules. (C) Molecular structures of the three pregnancy-related hormone molecules.

### Three Differentially Expressed Hormones

In this study, we identified three hormone-like metabolites that exhibited marked differences in expression, all of which are intimately linked to the physiological changes associated with pregnancy. These metabolites—Progesterone, 3-Deoxyestradiol, and Prostaglandin E1—demonstrated a significant upregulation in the vaginal secretions of pregnant sows, underscoring their pivotal role in gestational processes.

As illustrated in Table 1, all three hormones involved in pregnancy regulation exhibited notable sensitivity; however, 3-Deoxyestradiol distinguished itself by demonstrating superior performance, effectively differentiating between pregnant and non-pregnant samples with remarkable accuracy. Progesterone demonstrated a marginally lower Youden’s index compared to 3-Deoxyestradiol, whereas Prostaglandin E1 exhibited diminished specificity and Youden’s index, suggesting a compromised capacity to accurately distinguish non-pregnant samples.

**Table 1.** Diagnostic efficiency of ROC curve analysis of three hormone molecules.

| Metabolic molecules | Optimal threshold | Sensitivity | specificity | Youden Index |
| --- | --- | --- | --- | --- |
| Progesterone | 14825.35 | 0.87 | 0.90 | 0.77 |
| 3-Deoxyestradiol | 18109.84 | 0.93 | 1.00 | 0.93 |
| Prostaglandin E1 | 45421.28 | 0.87 | 0.70 | 0.57 |

### Machine learning analytics to build predictive models

In this investigation, two machine learning models—RF and SVM—were employed to elucidate the intricacies of untargeted metabolomics data, facilitating a comparative analysis of their classification efficacy for distinct sample sets. Following this, metabolites exhibiting statistically significant discrepancies between samples from pregnant and non-pregnant subjects were identified through a comprehensive integration of P-values, fold change (FC) metrics, and feature importance weights derived from both RF and SVM frameworks.

As illustrated in Figure 7, the RF model demonstrated an area under the ROC_AUC of 1.00 for both the training and testing datasets. The confusion matrix indicated an impeccable classification of the training set samples; however, a minor proportion of pregnant samples within the test set were inaccurately classified. The predictive probability boxplot demonstrated a markedly greater degree of predictive reliability for samples derived from pregnant individuals, contrasting sharply with the diminished reliability observed in samples from non-pregnant individuals. The learning curve illustrated that the model’s training set performance consistently exhibited a commendably high score. In contrast, the cross-validation score was initially low with a limited number of training samples. However, as the sample size surpassed 15, the cross-validation score progressively improved, resiliently converging towards the training set score.

**Fig. 7.**
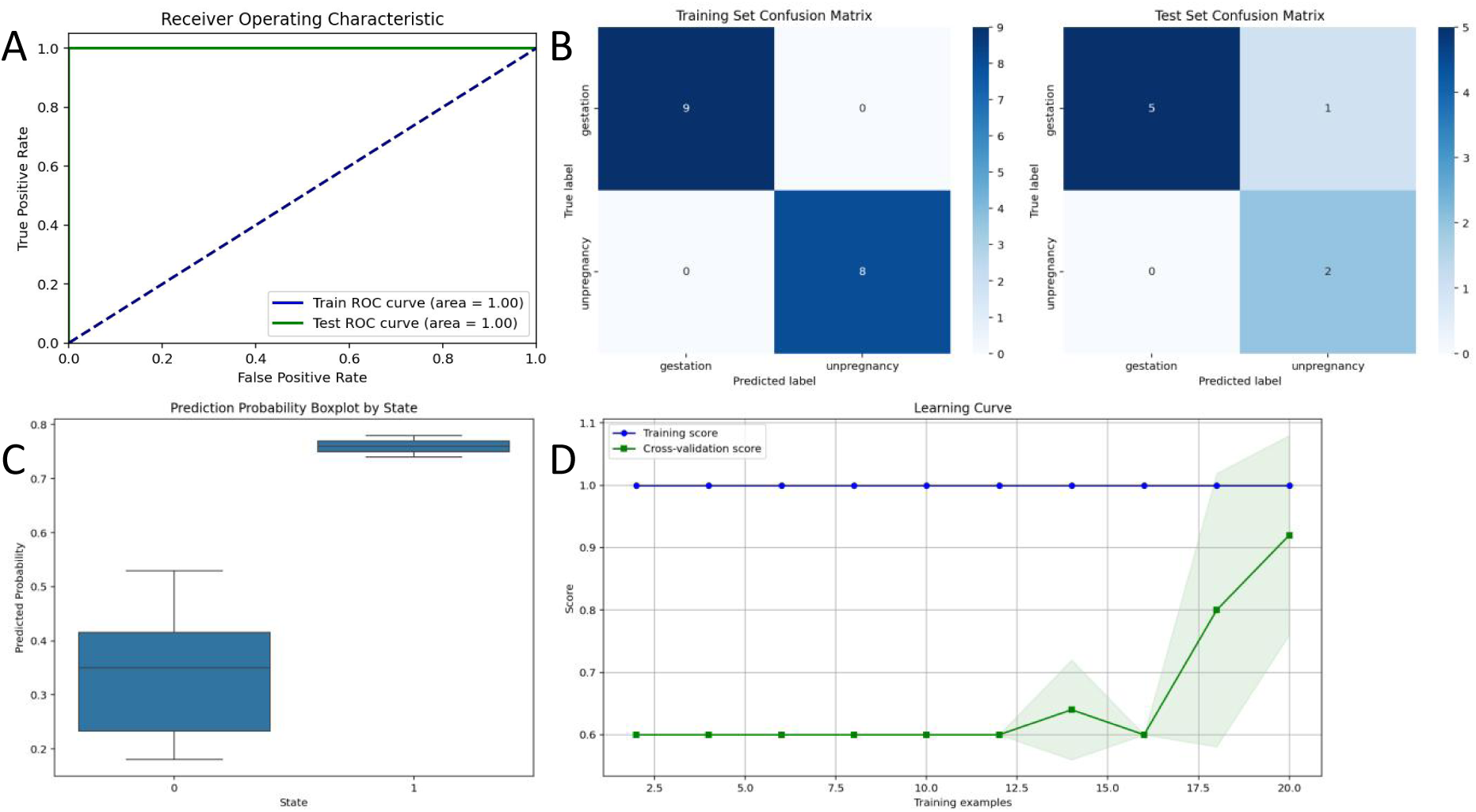
Random forest model parameter plots. (A) ROC plot. (B) Training and test set confusion matrices. (C) Predicted probability boxplot. (D) Learning curve.

As illustrated in Table 2, the RF model exhibited exemplary performance within the training dataset, achieving precision, recall, F1-score, and accuracy metrics, all of which attained a perfect value of 1.00. This outcome signifies an outstanding degree of data fitting. In contrast, the model’s performance on the test set exhibited a subtle decline, particularly evident in its capacity to predict non-pregnant samples, which demonstrated a comparatively lackluster efficacy.

**Table 2.** Random Forest model performance evaluation results.

|  |  | precision | recall | f1-score | support |
| --- | --- | --- | --- | --- | --- |
| training | pregnant | 1.00 | 1.00 | 1.00 | 9 |
|  | non-pregnant | 1.00 | 1.00 | 1.00 | 8 |
|  | accuracy |  |  | 1.00 | 17 |
|  | macro avg | 1.00 | 1.00 | 1.00 | 17 |
|  | weighted avg | 1.00 | 1.00 | 1.00 | 17 |
| testing | pregnant | 1.00 | 0.83 | 0.91 | 6 |
|  | non-pregnant | 0.67 | 1.00 | 0.80 | 2 |
|  | accuracy |  |  | 0.88 | 8 |
|  | macro avg | 0.83 | 0.92 | 0.85 | 8 |
|  | weighted avg | 0.92 | 0.88 | 0.88 | 8 |

As illustrated in Figure 8, in the SVM model, the ROC curves for both the training and test datasets attained an AUC value of 1.00. The confusion matrix revealed an impeccable classification, with no instances of misclassification present in either dataset. The predictive probability boxplot demonstrated a heightened degree of predictive reliability for pregnant samples, while exhibiting diminished reliability for non-pregnant samples. The analysis of the learning curve indicated that the model’s performance on the training set was persistently elevated, whereas the cross-validation scores exhibited a notable decline, particularly when the training sample size was limited. As the sample size exceeded a certain threshold, the cross-validation scores exhibited a progressive increase; however, they consistently remained considerably inferior to the scores obtained from the training set. This observation indicates a potential risk of overfitting.

**Fig. 8.**
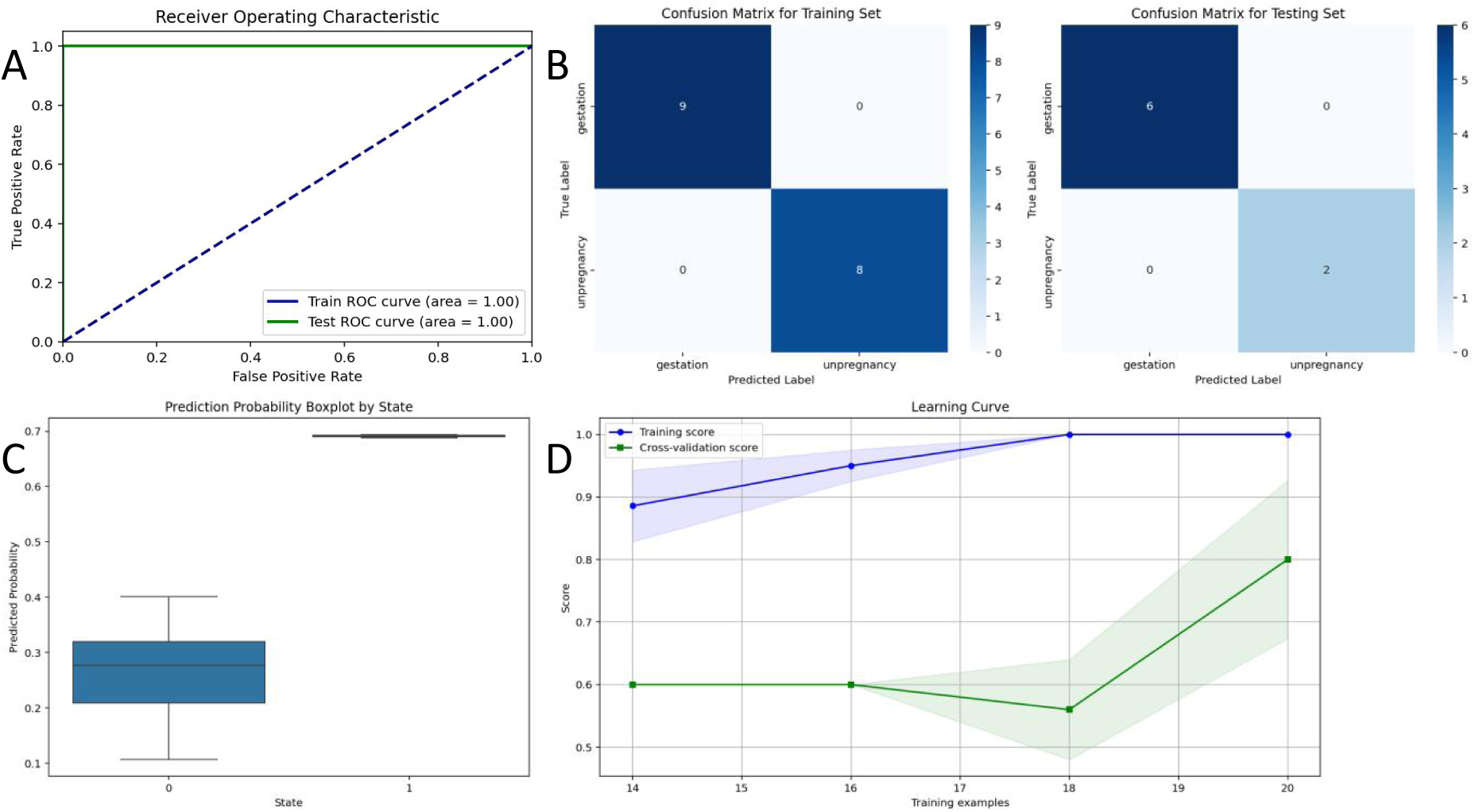
Support vector machine model parameter plots. (A) ROC plot. (B) Training and test set confusion matrices. (C) Predicted probability boxplot. (D) Learning curve.

As illustrated in Table 3, the SVM model exhibited markedly enhanced efficacy in the diagnosis of sow pregnancy through the analysis of vaginal secretions. It demonstrated remarkable predictive accuracy and reliability across both training and test datasets, adeptly distinguishing between samples from pregnant and non-pregnant sows based on the results of metabolomic profiling.

**Table 3.** Support vector machine model performance evaluation results.

|  |  | precision | recall | f1-score | support |
| --- | --- | --- | --- | --- | --- |
| training | pregnant | 1.00 | 1.00 | 1.00 | 9 |
|  | non-pregnant | 1.00 | 1.00 | 1.00 | 8 |
|  | accuracy |  |  | 1.00 | 17 |
|  | macro avg | 1.00 | 1.00 | 1.00 | 17 |
|  | weighted avg | 1.00 | 1.00 | 1.00 | 17 |
| testing | pregnant | 1.00 | 1.00 | 1.00 | 6 |
|  | non-pregnant | 1.00 | 1.00 | 1.00 | 2 |
|  | accuracy |  |  | 1.00 | 8 |
|  | macro avg | 1.00 | 1.00 | 1.00 | 8 |
|  | weighted avg | 1.00 | 1.00 | 1.00 | 8 |

### Significance of difference analysis combined with machine learning to screen target metabolites

As illustrated in Figure 9, this investigation identified 510 metabolic molecules exhibiting significant differences in their concentrations, alongside notable up- or down-regulation between pregnant and non-pregnant samples. Additionally, 123 metabolic molecules were identified that exerted influence within the RF model, while 127 metabolic molecules demonstrated an impact weight exceeding 0.3% on the decision boundary within the SVM model. The Venn diagrams derived from these three screening methodologies revealed three overlapping metabolite molecules: Indolepropionic acid (POS2295), cis-Aconitate (NEG2539), and 4-p-Coumaroylquinic acid (NEG13945).

**Fig. 9.**
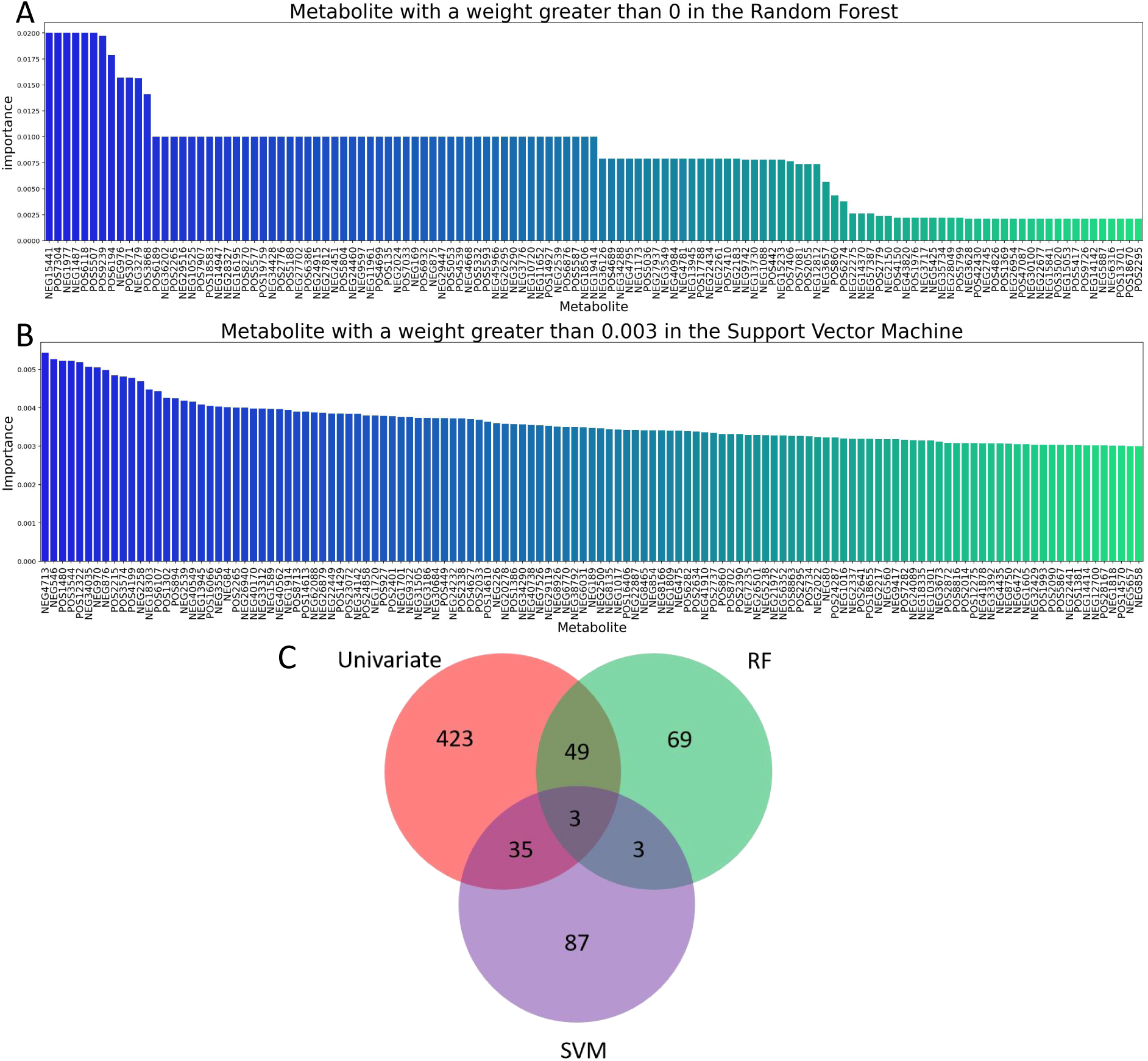
The three methods work together to screen for differentially metabolized molecules. (A) Metabolic molecules with influence on the model in the RF model. (B) Metabolic molecules with more than 0.3% influence weight on the model in the SVM model. (C) Wayne plots of metabolic molecules with significant differences in expression, RF influence weights, and SVM influence weights that are high.

As illustrated in Figure 10, the ROC curves reveal commendably high AUC values for each of the three metabolic molecules examined. Indolepropionic acid exhibits an AUC of 0.90, followed closely by cis-Aconitate at 0.89, and 4-p-Coumaroylquinic acid with a somewhat lower AUC of 0.79. Notably, the expression of Indolepropionic acid is significantly heightened in pregnant samples when compared to their non-pregnant counterparts, whereas expressions of both cis-Aconitate and 4-p-Coumaroylquinic acid are markedly diminished. The three metabolic molecules demonstrated potential as diagnostic marker metabolites for detecting pregnancy in sows through the analysis of vaginal secretions.

**Fig. 10.**
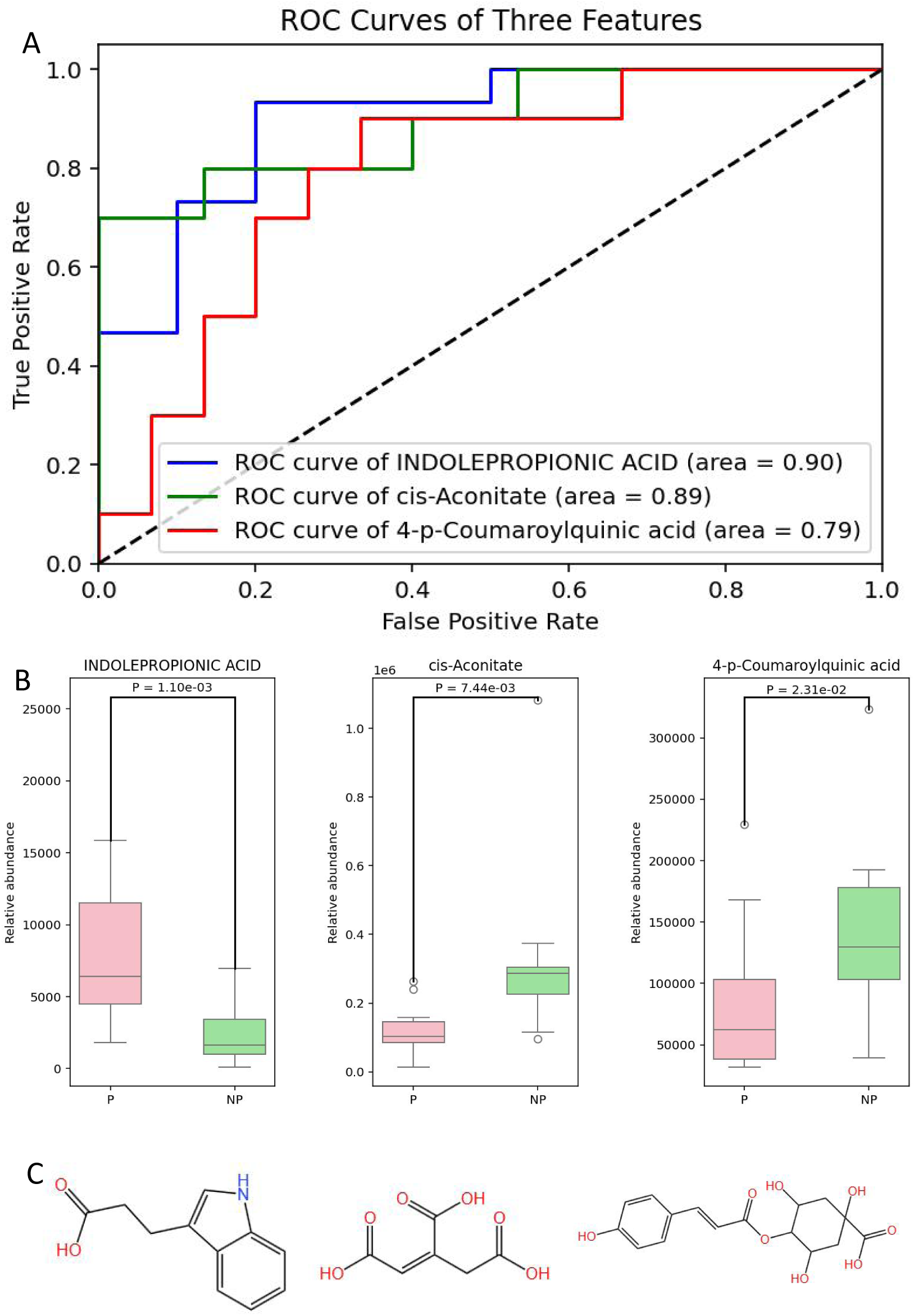
Three target metabolic molecules. (A) ROC curves of the three target molecules (B) Differences in expression of the three target molecules in different groups of samples (C) Structures of the three target molecules.

As illustrated in Table 4, the three targeted metabolic molecules exhibit varying efficacies in diagnosing pregnancy in sows. Among these, Indolepropionic acid demonstrates remarkable sensitivity and specificity, coupled with an elevated Youden index, establishing it as an exemplary diagnostic biomarker for pregnancy. Conversely, while cis-Aconitate presents high specificity, its sensitivity is comparatively lower, indicating that it is more appropriately utilized for excluding nulliparous pregnancies rather than for affirming gestation. In contrast, 4-p-Coumaroylquinic acid displays greater sensitivity but diminished specificity, resulting in a propensity for misclassification.

**Table 4.**
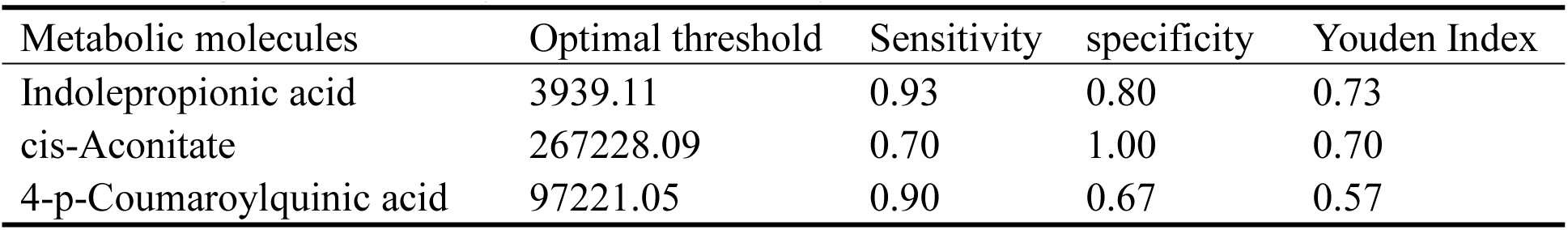
Diagnostic efficiency of ROC curve analysis for three classes of metabolic molecules.

## DISCUSSION

This study introduces an innovative methodology for utilizing vaginal secretions in the early diagnosis of pregnancy in sows. In contrast to other bodily fluids, such as urine and blood, vaginal secretions present unique advantages due to their non-invasive sampling technique and comparatively stable physicochemical characteristics. Furthermore, owing to the anatomical proximity of the vaginal tract to the sow’s uterus, it demonstrates heightened sensitivity to the physiological alterations occurring within the uterus. This proximity renders it a particularly suitable medium for non-invasive surveillance of uterine status.

Nonetheless, the vagina’s numerous glands and diverse microbial communities introduce various confounding variables that influence the physicochemical characteristics of vaginal secretions. This finding suggests that employing vaginal secretions to identify specific physiological states in sows may be influenced by uncontrollable variables. In this investigation, PCA of vaginal secretion samples revealed a modest trend of separation between pregnant and non-pregnant groups; however, this distinction lacked statistical significance. Consequently, this underscores the potential risks of misclassification when relying exclusively on PCA for this purpose. In contrast, the PLS-DA analysis revealed a more definitive pattern of separation, likely due to the fact that PCA emphasizes data variance exclusively. This limitation becomes particularly salient considering that the metabolomics of vaginal secretions encompass high-variance features that may not be relevant to pregnancy status. These characteristics obscure nuanced linear disparities within the data, which PLS-DA adeptly resolves through supervised learning by extracting components of variance associated with pregnancy.

In the volcano plot delineating differential metabolites, a total of 323 metabolites exhibited significant up-regulation, whereas 187 demonstrated notable down-regulation. Significantly, three key hormones involved in pregnancy regulation — Progesterone, 3-Deoxyestradiol, and Prostaglandin E1 — demonstrated substantially elevated expression levels in the vaginal secretions of pregnant sows in contrast to their non-pregnant counterparts. This observation elucidates notable physicochemical disparities in vaginal secretions between the two groups, likely arising from the physiological alterations associated with pregnancy [12–14].

Progesterone has been widely employed as a biomarker for diagnosing pregnancy across a diverse array of mammalian species, including humans, pigs, buffalo, goats, and sheep. For example, Aly Karen and colleagues utilized assays for progesterone and pregnancy-associated glycoproteins to facilitate the early detection of pregnancy in sheep [15]. Meanwhile, J.A. Pennington and associates assessed progesterone levels in milk as a diagnostic tool for pregnancy in dairy goats [16]. Furthermore, Perera B.M. and team employed plasma progesterone concentrations for the early diagnosis of pregnancy in buffaloes [17]. In contrast,

3-Deoxyestradiol and Prostaglandin E1 have not been documented in the context of pregnancy diagnosis. While 3-Deoxyestradiol is hypothesized to be a metabolite of estradiol—though this remains unverified—its pronounced differential expression in vaginal secretions of sows indicates its promising potential as a diagnostic biomarker for pregnancy.

Prostaglandin E1, an endogenous metabolite integral to processes such as vasodilation, inhibition of platelet aggregation, and the regulation of reproductive functions, exhibited a marked upregulation in the vaginal secretions of pregnant sows. Nevertheless, its comparatively modest area under the ROC curve and Youden’s index imply that its expression may remain elevated even in non-pregnant specimens, thereby constraining its utility as a solitary diagnostic biomarker [18].

The KEGG pathway enrichment analysis, complemented by two-tier KEGG pathway classification visualizations, revealed a pronounced enrichment of differential metabolites among the metabolic pathways of vaginal secretions from pregnant sows compared to their non-pregnant counterparts. Notably, nucleotide metabolism emerged as one of the most significant areas of focus. This suggests that the metabolic processes of nucleotides experience significant transformations within the vaginal milieu throughout the course of pregnancy. Throughout gestation, the reproductive system of the sow undergoes significant activation, marked by an upsurge in cellular proliferation, differentiation, and tissue repair processes. Given that nucleotides are fundamental constituents for the synthesis of both DNA and RNA, the activation of nucleotide metabolic pathways is vital in meeting the heightened demands of extensive cellular proliferation and tissue regeneration. This physiological process subsequently results in the accumulation of associated metabolites within vaginal secretions [19]. Furthermore, the hormonal fluctuations associated with pregnancy—most notably the increased concentrations of progesterone and estrogen—are instrumental in modulating intracellular gene expression as well as metabolic pathways. These hormones stimulate the biosynthesis and degradation of nucleotides by regulating the expression of enzymes that play a pivotal role in nucleotide metabolism[20,21]. This regulatory mechanism further elucidates the metabolic transformations that occur throughout gestation.

Furthermore, the immune system of gestating sows experiences a nuanced adaptation, intricately designed to safeguard the developing fetus. Given that the vagina functions as the primary barrier against pathogenic infiltration, it is plausible that modifications in both the activity and quantity of immune cells may transpire. The expansion of immune cell populations and the synthesis of immune molecules are contingent upon the participation of nucleotides, which in turn amplifies nucleotide metabolism and elevates the levels of nucleotide-associated metabolites in vaginal secretions [22].

In the development and validation of machine learning models, both models exhibited remarkable fitting and predictive capabilities for untargeted metabolomics data, thereby affirming the reliability of the metabolomic findings. Three pivotal metabolic molecules—Indolepropionic acid, cis-Aconitate, and 4-p-Coumaroylquinic acid—were meticulously screened, predicated on their expression significance and feature weights derived from machine learning models. This analysis reveals a pronounced and significant disparity in their stable expression levels within vaginal secretions of pregnant versus non-pregnant sows.

Indolepropionic acid, an indole derivative synthesized by gut microbiota during the metabolic processing of tryptophan, plays multifaceted physiological roles in animals. These roles encompass the preservation of intestinal microbial homeostasis, augmentation of intestinal barrier integrity, enhancement of metabolic regulation, modulation of immune responses, neuroprotective effects, and the regulation of neurotransmitter metabolism [23]. Throughout gestation, fluctuations in progesterone levels in sows play a crucial role in mitigating intestinal inflammation, fostering the proliferation of probiotics that produce indole-3-propionic acid, and curbing the overgrowth of pathogenic bacteria. This intricate interplay ultimately enhances the expression levels of Indolepropionic acid[24]. Simultaneously, estrogens have the potential to upregulate intestinal tryptophan transporter proteins, such as SLC7A5, thereby augmenting the availability of tryptophan for the gut microbiota and enhancing the precursor supply for indole-3-propionic acid biosynthesis [25]. Moreover, placental cytokines, such as interleukin-10 (IL-10) and tumor necrosis factor-alpha (TNF-α), may influence the metabolic activity of gut microorganisms through systemic circulation. This modulation potentially enhances the production of indole-3-propionic acid and facilitates its diffusion into the bloodstream, ultimately reaching the reproductive tract [26,27].

Cis-aconitate, a pivotal intermediate in the intracellular tricarboxylic acid (TCA) cycle, assumes a critical role in the metabolic processes and physiological functions of animal systems. As a crucial juncture in the energy metabolism framework of the TCA cycle, its upstream and downstream metabolites—such as citrate and isocitrate—contribute to essential pathways, including fatty acid biosynthesis, gluconeogenesis, and amino acid metabolism [28,29]. Concurrently, aconitase—the enzyme responsible for the catalysis of cis-aconitate formation—exists as two principal isoforms: mitochondrial and cytoplasmic. The function of cytosolic aconitase is intricately associated with the regulation of iron metabolism [30–32].

In pregnant sows, the demands of energy metabolism transition from sustaining maternal homeostasis to facilitating embryonic development, placental formation, and the maturation of mammary glands. This condition enhances the metabolic demands of reproductive organs, such as the uterus and placenta, thereby promoting the preferential allocation of systemic TCA cycle intermediates, notably cis-aconitate, to the feto-placental unit. Simultaneously, there is a corresponding decline in cis-aconitate concentrations within vaginal secretions [33]. The gestational phase engenders a glucose partitioning phenomenon reminiscent of “gestational diabetes,” whereby glucose is preferentially allocated to the uterus and mammary glands, resulting in a diminished uptake in the vaginal epithelium. This alteration leads to a decrease in cis-aconitate production through the pathways of glycolysis and the TCA cycle [34].

The secretion of progesterone, driven by pregnancy and originating from both the corpus luteum and placenta, further modulates the levels of cis-aconitate. To impede uterine contractions and sustain pregnancy, progesterone may diminish the proliferation and metabolic activity of vaginal epithelial cells, simultaneously inhibiting the expression or activity of aconitase. This interplay consequently leads to a reduction in the concentration of cis-aconitate within vaginal secretions [35]. Fluctuations in estrogen levels are also significant: following pregnancy, estrogen concentrations are generally reduced compared to those in estrous sows, which leads to a depletion of glycogen reserves in vaginal epithelial cells. This diminishes the flux of pyruvate into the mitochondrial TCA cycles through glycolysis, thereby leading to a reduction in the synthesis of cis-aconitate [36].

4-p-Coumaroylquinic acid, a naturally occurring phenolic acid prevalent in various plant species, is predominantly found in animals as a metabolic byproduct following dietary consumption [37]. Although its functions in animals as potential drug targets are not yet fully elucidated, research has substantiated its antioxidant, anti-inflammatory, and metabolic regulatory properties [38]. As a phenolic acid, it demonstrates antioxidant activity through the scavenging of free radicals, the inhibition of oxidative enzymes, and the augmentation of overall antioxidant capacity. Simultaneously, it attenuates the expression of inflammatory factors (such as TNF-α, IL-6, and IL-1β), downregulates the activation of the NF-κB signaling pathway, and diminishes the activity of cyclooxygenase-mediated inflammatory enzymes.

In pregnant sows, the interplay of hormonal regulation, physiological metabolic adaptations, and the remodeling of the reproductive tract microenvironment significantly influences the composition of vaginal secretions [39]. For example, the concentrations of estrogen and progesterone in both pregnant and non-pregnant sows may influence metabolic processes: estrogen stimulates the activity of hepatic cytochrome P450 enzymes, which in turn induces the expression of phenolic acid-metabolizing enzymes within vaginal mucosal cells. This upregulation accelerates the metabolism and excretion of these compounds, ultimately resulting in diminished concentrations in both the circulatory system and reproductive tract [40]. Conversely, progesterone may inhibit the intestinal absorption of dietary phytophenolic acids, subsequently influencing their levels of secretion in the vaginal milieu. Throughout gestation, the vaginal epithelium may prioritize the metabolism of glycogen in response to hormonal influences, which consequently leads to a reduction in the synthesis or secretion of phenolic acids [41,42].

Conventional methods for diagnosing pregnancy in sows frequently encounter challenges such as the induction of maternal stress, operational complexities, and suboptimal timing [43,44]. Ultrasonography, the predominant technique employed in swine husbandry, necessitates an interval of 23 to 25 days following mating to achieve reliable detection of pregnancy. Nevertheless, given that non-pregnant sows display a 21-day estrous cycle, this methodology frequently results in the oversight of optimal re-mating opportunities. In this study, the analysis of vaginal secretions for pregnancy diagnosis at 18 days post-mating not only alleviated stress in gestating sows but also enhanced the effective detection window by 5 to 8 days in comparison to traditional ultrasonography. This methodology enables farms to discern the most advantageous re-mating opportunities for sows that are not currently pregnant.

## CONCLUSION

This study adeptly identified 510 distinctively expressed metabolites in the vaginal secretions of both pregnant and non-pregnant sows. Three crucial hormones involved in pregnancy regulation—progesterone, 3-deoxyestradiol, and prostaglandin E1—exhibited a significant upregulation in the vaginal secretions of pregnant sows. Furthermore, machine learning modeling identified three metabolic molecules exhibiting significant differential expression: indolepropionic acid, cis-aconitate, and 4-p-coumaroylquinic acid. These findings substantiate the application of vaginal secretions as an effective metabolic matrix for the early diagnosis of pregnancy in sows, thus establishing a crucial scientific foundation for the advancement of non-invasive pregnancy diagnostic techniques within swine production.

## AUTHOR CONTRIBUTIONS

T.H provided the research idea.

Y.F and T.H designed the study.

Y.F, R.G, and Yun.Z performed the sample collection and analyzed the data.

Y.F, P.G, W.Y, Yongsheng.Z, Y.R, Z.A, K.Y, H.L, and T.H provided reagents/materials/analysis tools/technical support.

Y.F and M.L wrote the manuscript.

M.L, Q.L, and T.H revised the manuscript.

All authors read and approved the final version of the manuscript.

## ACKNOWLEDGMENTS

The authors would like to express their gratitude to Shanghai Bioprofile for the technical support and assistance in data analysis.

## DATA AVAILABILITY

Upon reasonable request, the datasets of this study can be available from the corresponding author.

## ETHICS APPROVAL

This animal study was reviewed and approved by the Bioethics Committee of Shihezi University. All experimental procedures were conducted strictly in accordance with the local administrative regulations on animal experiments and the institutional ethical requirements for animal research. Prior to the initiation of the study, written informed consent was formally obtained from the owners of the animals involved, confirming their permission for the animals to participate in this research.

## DECLARATION OF GENERATIVE AI

During the preparation of this work, DouBao was used for the purpose of translate. After using this tool, the manuscript was reviewed and edited as needed and authors will assume full responsibility for the publication.

